# Designing β-hairpin peptide macrocycles for antibiotic potential

**DOI:** 10.1101/2022.06.21.497034

**Authors:** Justin R. Randall, Cory D. DuPai, T. Jeffrey Cole, Gillian Davidson, Kyra E. Groover, Claus O. Wilke, Bryan W. Davies

**Affiliations:** Department of Molecular Biosciences, University of Texas at Austin; Austin, TX, USA; Department of Integrative Biology, University of Texas at Austin; Austin, Texas, USA

## Abstract

Peptide macrocycles are a rapidly emerging new class of therapeutic, yet the design of their structure and activity remains challenging. This is especially true for those with β-hairpin structure due to weak folding properties and a propensity for aggregation. Here we use proteomic analysis and common antimicrobial features to design a large peptide library with macrocyclic β-hairpin structure. Using an activity-driven high-throughput screen we identify dozens of peptides killing bacteria through selective membrane disruption and analyze their biochemical features via machine learning. Active peptides contain a unique constrained structure and are highly enriched for cationic charge with arginine in their turn region. Our results provide a synthetic strategy for structured macrocyclic peptide design and discovery, while also elucidating characteristics important for β-hairpin antimicrobial peptide activity.

**Brief Summary:** We design, screen, and computationally analyze a synthetic macrocyclic β-hairpin peptide library for antibiotic potential.

## Introduction

The stability and broad functionality of macrocyclic peptides makes them a promising area for drug development; however, there are few well-characterized strategies for their identification. Current *de novo* design remains challenging, especially for those with β-hairpin structure (*1*–*4*). They often lack sufficient inter-strand interactions to form stable conformations and their β-strands promote association which can lead to aggregation in solution (*3*–*5*).

Antibiotics provide an excellent example of macrocyclic peptide drug value and new discovery strategies are urgently needed (*6*). Macrocyclic β-hairpin antimicrobial peptides (β-AMPs) have recently gained popularity, with two such antibiotics having proceeded into clinical evaluation (*7*–*9*). These β-AMPs act primarily through bacterial membrane permeabilization, allowing them to overcome most mechanisms of bacterial drug resistance and also access and inhibit essential processes within the Gram-negative cell envelope (*10*, *11*); however, known examples are exceedingly rare. This makes it is difficult to understand how β-AMP sequence determines structure and function (*12*–*14*). Established strategies for *de novo* β-hairpin peptide design also limit the use of amino acids important for antibacterial activity and include residues that increase mammalian cell toxicity, which is detrimental for therapeutic development (*1*, *12*).

Here we describe the design, screening, and analysis of a synthetic macrocyclic β-hairpin peptide library using new strategies and technologies. Our results expand our understanding of β-AMP sequence-activity relationships and provides a new route for their design and discovery.

## Results

### Design of a synthetic macrocyclic β-hairpin peptide library

Our design scheme leveraged our recently completed systematic analysis of over 49,000 β-hairpin motifs in the Protein Data Bank. This analysis identified position specific amino acid preferences in the strand and turn regions (*15*). Using this information, we designed two ribosomally encoded twenty amino acid cyclic β-hairpin peptide libraries (**Fig. 1AB** and **fig. S1**).

**Figure 1.**
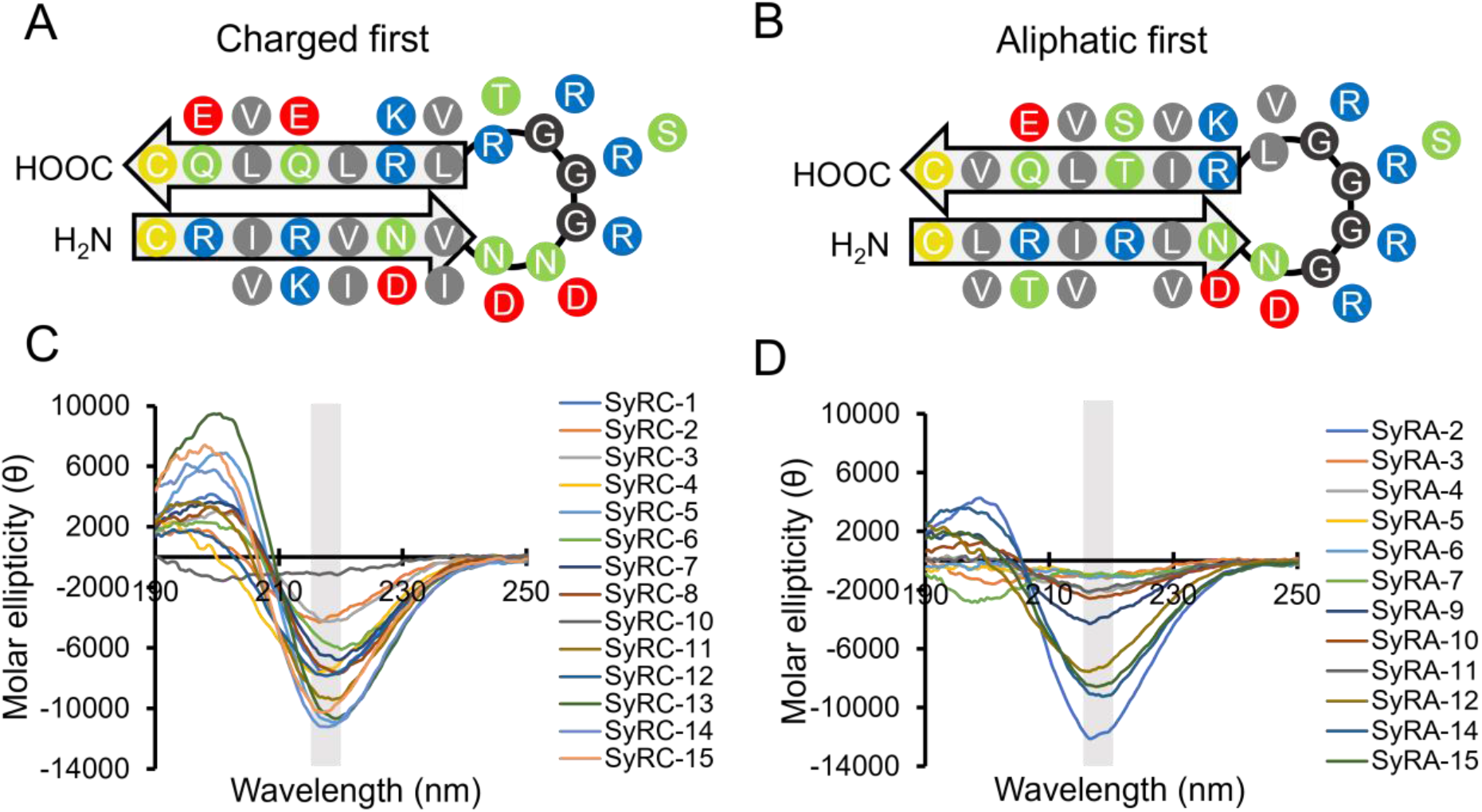
SynCH peptides show β-hairpin secondary structure. Diagrams showing the designed structure and residue position of the charged first (**A**) and aliphatic first (**B**) SynCH libraries. Residues are color coded by side chain (yellow = cysteine, green = polar, gray = aliphatic, blue = positive, red = negative). (**C**, **D**) Circular dichroism spectra of randomly selected charged first (SyRC) and aliphatic first (SyRA) SynCH peptides. A single minimum between 215 and 220 nm (gray box) is characteristic of β-hairpin secondary structure. Each spectrum is the mean of three technical replicates with background subtracted.

These libraries were intentionally designed to facilitate the development of stable antiparallel β-strands with amphipathic faces generated via the periodic alternation of aliphatic and charged/polar residues. We identified and applied a preference for glycine, asparagine, and aspartic acid in or near the turn regions (*15*–*17*) and excluded proline in contrast to canonical β-turn design (*18*–*21*). We also excluded aromatic residues because they promote mammalian toxicity despite their positive effects on β-hairpin structure (*22*). Cysteine was encoded at the amino and carboxyl termini to potentiate cyclization via a disulfide bridge and prevent fraying. Finally, we allowed select polar positions and the loop region to encode for positive residues to promote solubility and to mimic a trend we observed in natural β-AMPs. Using codon variation, we created and combined two synthetic, macrocyclic β-hairpin (SynCH) libraries based on our design scheme, one beginning with a charged residue and the other with an aliphatic residue. This allows different amphipathic characters to occupy different faces relative to the loop region. In total, this pooled library encompassed 196,608 unique peptide sequences possessing a variety of physiochemical properties. (**Fig. 1AB** and **fig. S1**).

### SynCH peptides spontaneously fold into macrocyclic β-hairpins

We examined the structure of thirty SynCH peptides at random from the charged first (SyRC) and aliphatic first (SyRA) libraries spanning a range of charge (−0.12 to 4.87) and grand average of hydropathicity (GRAVY) score (−0.72 to 0.41) (**Table 1**). One SyRC peptide and three SyRA peptides could not be synthesized. We first used circular dichroism spectroscopy (CD) to determine the secondary structure of these chemically synthesized peptides (**Fig. 1CD, Table 1**). Remarkably, 84.6% of all peptides had a CD spectrum with a single ellipticity minimum between 215 and 220 nm, indicative of antiparallel β-sheet secondary structure (*23*).

**Table 1.**
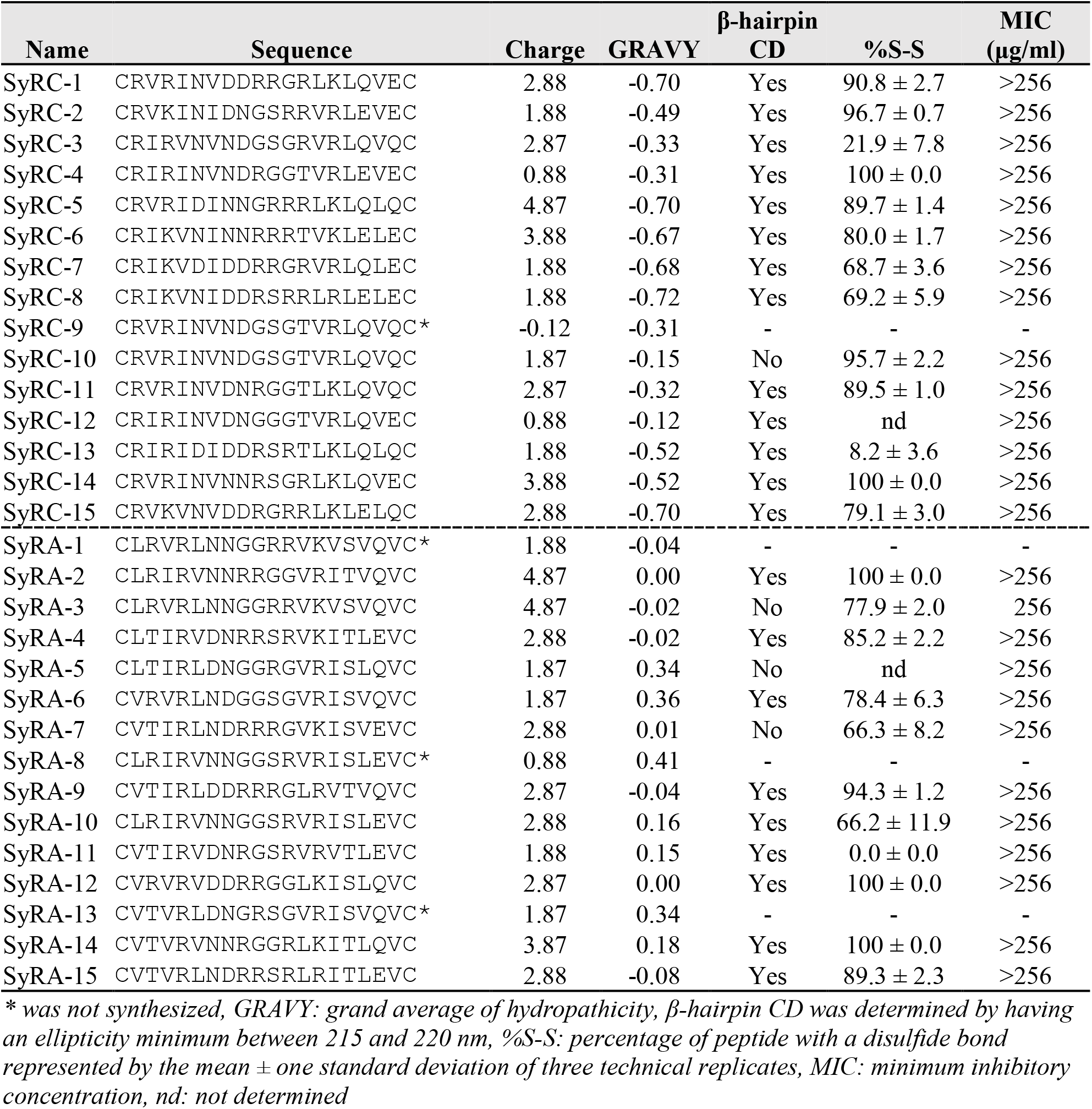
Properties of randomly selected peptides from the SynCH library

This is highly uncommon for peptides in aqueous solution. Most require a target interaction to form a β-hairpin or other secondary structure (*24*, *25*).

Next, we performed high resolution LC/MS on each peptide to determine whether an intramolecular disulfide bond was present (**Table 1, fig. S2**). All but three of the randomly selected SynCH peptides examined had a disulfide bond present in the majority of their molecular population. While sequences from the aliphatic first library were less likely to show β-sheet secondary structure (**Fig. 1CD**), they were slightly more likely to have a majority of their molecular population cyclized (**Table 1**). This data together suggests that ~72% of our SynCH peptide library forms a stable β-hairpin secondary structure and are cyclized through an intramolecular disulfide bond in solution.

### Identification of SynCH peptides with antibiotic potential using SLAY

Our randomly selected SynCH peptides were not inherently antibacterial (**Table 1**), so we decided to use a high-throughput genetic platform developed in our lab called surface localized antimicrobial display (SLAY) (*26*, *27*) to screen for antibacterial activity. SLAY functions through the inducible display of a plasmid encoded peptide library on the Gram-negative bacterial cell surface. Next-generation sequencing is then used to generate a log_2_-fold change in peptide sequence read counts between induced and uninduced bacterial cultures (**Fig. 2A**).

**Figure 2.**
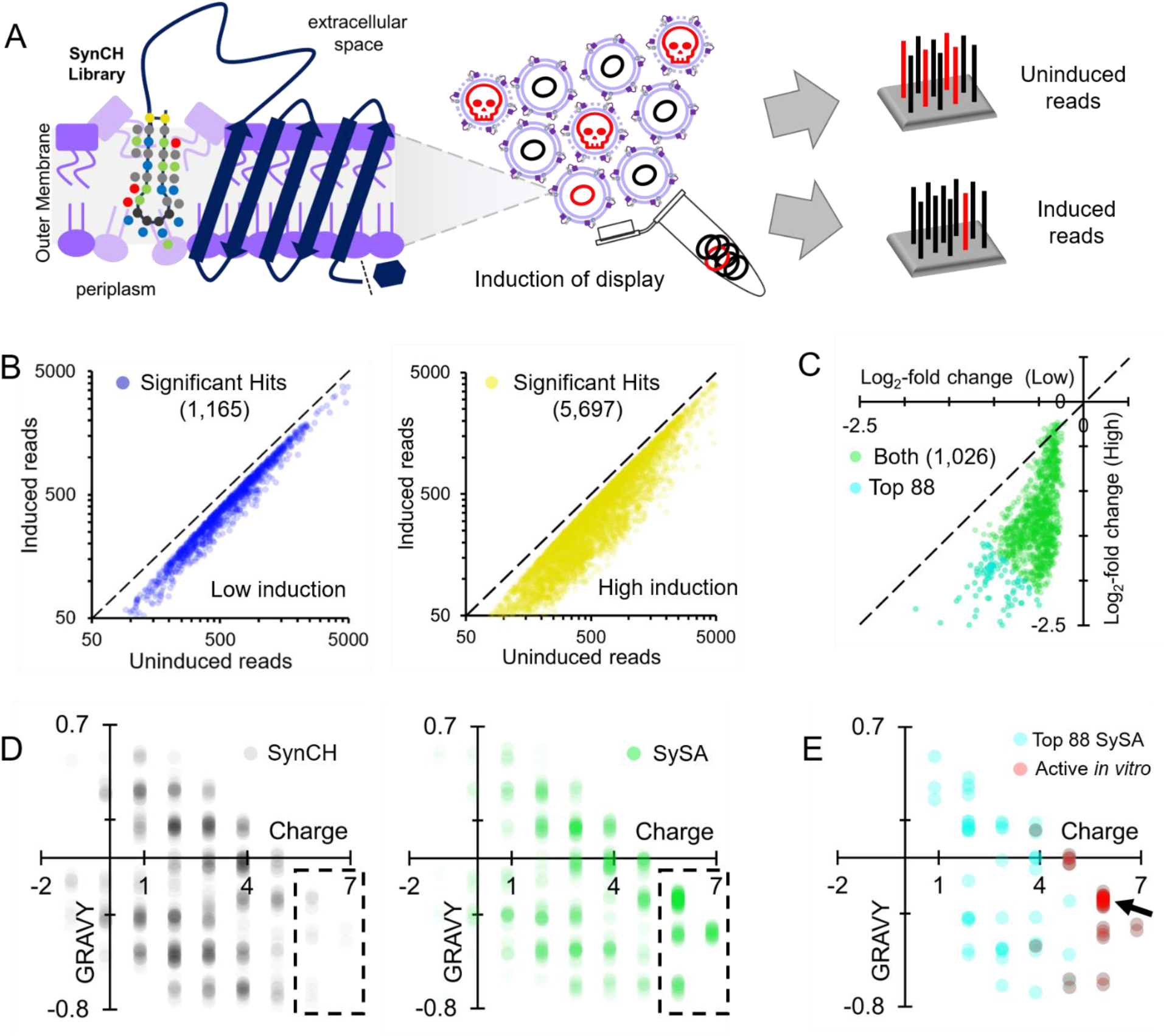
SLAY identifies SynCH peptides with antibacterial potential. (**A**) Workflow of a surface localized antimicrobial display (SLAY) screen. (**B**) Density scatter plot for SynCH peptides with a significant, negative log_2_-fold change (p < 0.05) from SLAY induced at low (left) and high (right) concentration. (**C**) Density scatter plot of each SynCH peptide with a significant, negative log_2_-fold change in reads (p < 0.05) under both induction conditions (SySA). (**D**) Density scatter plots for a randomized subset of the SynCH library (left) and SySA (right). (**E**) Same plot but with the top 88 SySA peptides based on average log_2_-fold change. Those which were verified to have in vitro activity are in red, 18/36 are from a single grouping (arrow).

We performed SLAY on our SynCH library at low (15 μM IPTG) and high (100 μM IPTG) induction concentrations to mimic treating bacteria with two different concentrations of peptide. The low induction (15 μM IPTG) condition identified 1,165 such peptides, referred to as “Hits” (**Fig. 2B, left**). The high induction (100 μM IPTG) condition identified significantly more “Hits” (5,697) (**Fig 2B, right**). Next, we plotted the log_2_-fold change of the hits from both induction concentrations against one another (**Fig. 2C**). 1,026 peptides, or 2.49% of the total number of peptides screened (~41,000), were identified as a “Hit” under both conditions. This group of 1,026 peptides are referred to as the SynCH SLAY Active group, or “SySA” going forward (**Fig. 2C, green**). The most promising 88 hits are also highlighted (**Fig. 2C**, cyan).

To see if there was enrichment of SynCH peptides with a certain charge or hydrophobicity in the SySA group, we plotted the distribution of charge versus GRAVY score for an equal number of randomly selected SynCH peptides and compared them to the SySA group (**Fig. 2D**). SySA distribution was highly enriched at charges greater than 5.5 relative to the SynCH library, suggesting that cationic charge is important for their antibacterial potential. The top 88 SySA peptides were further enriched toward cationic charge (**Fig. 2E, cyan**). Interestingly, 18/36 SySA peptides with verified *in vitro* activity (detailed below) were within a single distribution grouping (charge ~5.87, GRAVY −0.2 to −0.3) (**Fig. 2E, red, arrow**).

### SySA peptides selectively disrupt bacterial membranes

We chose to chemically synthesize the top 88 SySA peptides as determined by lowest average log2-fold change for further biochemical characterization (**Fig. 2CE cyan**) along with two natural β-AMPs for comparison: Protegrin-1 and Thanatin. We began by determining the minimum inhibitory concentration (MIC) for each SySA peptide against our screening strain, *E. coli* W3110 in both Mueller-Hinton broth (MH) and the tissue culture media RPMI 1640 (RPMI). MH broth is a standard for antibacterial testing, while RPMI medium better represents salt and buffer conditions found in the body. 44.4% (36/81) of the SySA peptides examined were active *in vitro* in both media with MICs ranging from 4 to 256 μg/ml (**Table 2**). Interestingly, the vast majority of active peptides were from the aliphatic first library. All active SySA peptides had a charge greater than 3.8 with 78% having a charge greater than 5.8. Interestingly, 18 of the 36 peptides had a charge of 5.87 and a GRAVY score between −0.2 and −0.3 (**Fig. 2E,** arrow; **Table 2,** bolded). This grouping contained many of the most potent SySA peptides, so we chose five (MICs 4-32 μg/ml) to investigate further (**Fig. 3A**). Overall, this set of peptides was less potent than Protegrin-1 and Thanatin, whose MICs ranged from 0.5-4 μg/ml (**Fig. 3A**). This is not surprising considering our SySA peptides have not been optimized for antibacterial activity, while natural β-AMPs have evolved their activity over millennia.

**Figure 3.**
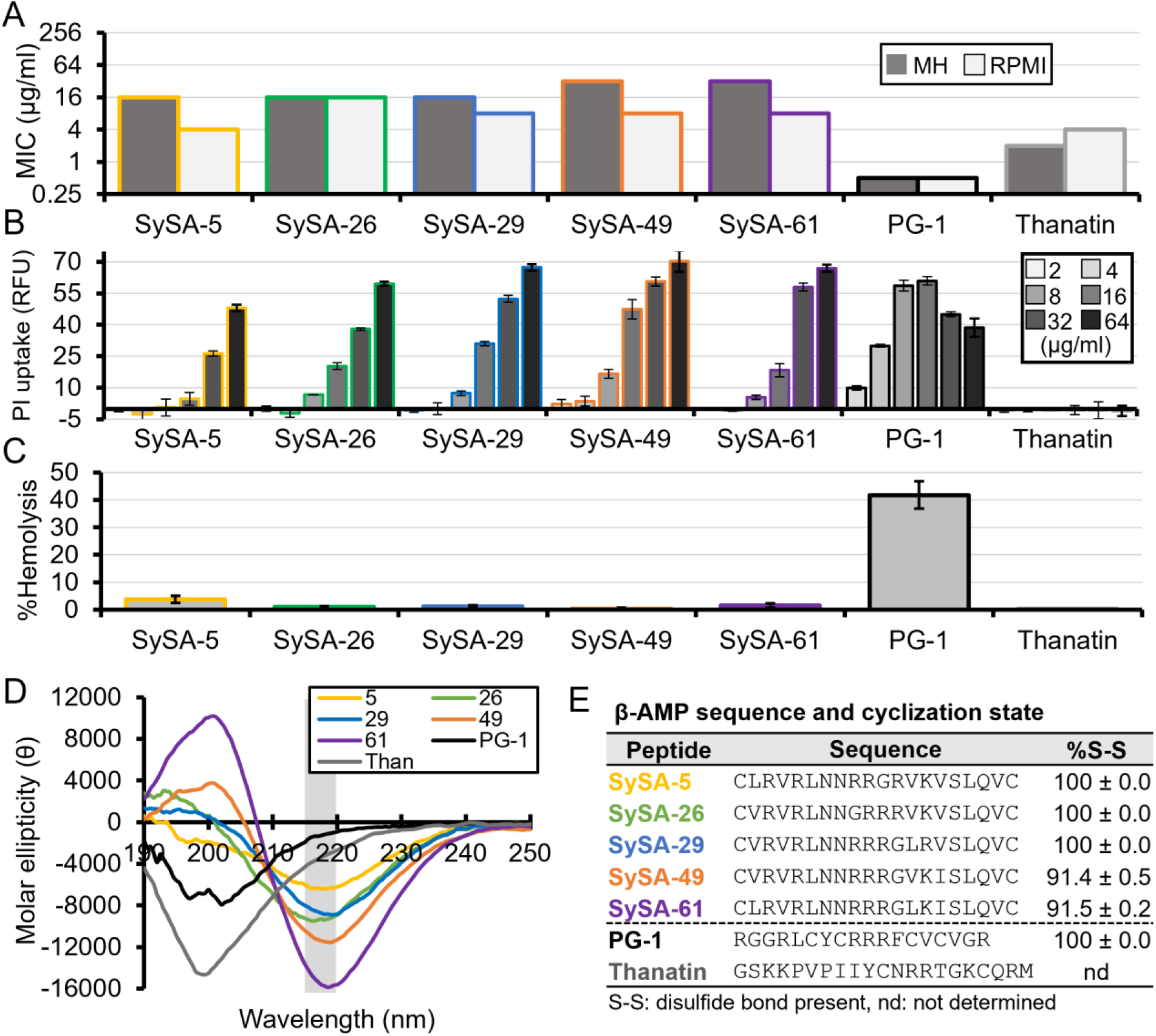
SySA peptide antibiotics demonstrate membrane specificity, have cyclic β-hairpin structure. (**A**) Minimum inhibitory concentration of select SySA peptides, Protegrin-1 (PG-1) and Thanatin against E. coli W3110 in Mueller-Hinton (MH) and RPMI1640 (RPMI) media. Each bar represents the median MIC (n=3) (**B**) The %hemolysis relative treatment with 1% Triton-X100. Bars represent the mean; error bars are one standard deviation (n=3) (**C**) Propidium iodide (PI) uptake of E. coli W3110 cells treated with peptide. Bars represent the median and error bars one standard deviation (n=3) (**D**) Circular dichroism spectra of peptides. A single minimum between 215 and 220 nm (gray box) is characteristic of β-hairpin structure. Each spectrum is the mean of three technical replicates with background subtracted. (**E**) Table of each peptides sequence and the percentage of the molecular population with an intramolecular disulfide bond. Error is one standard deviation of technical replicates (n=3).

**Table 2.**
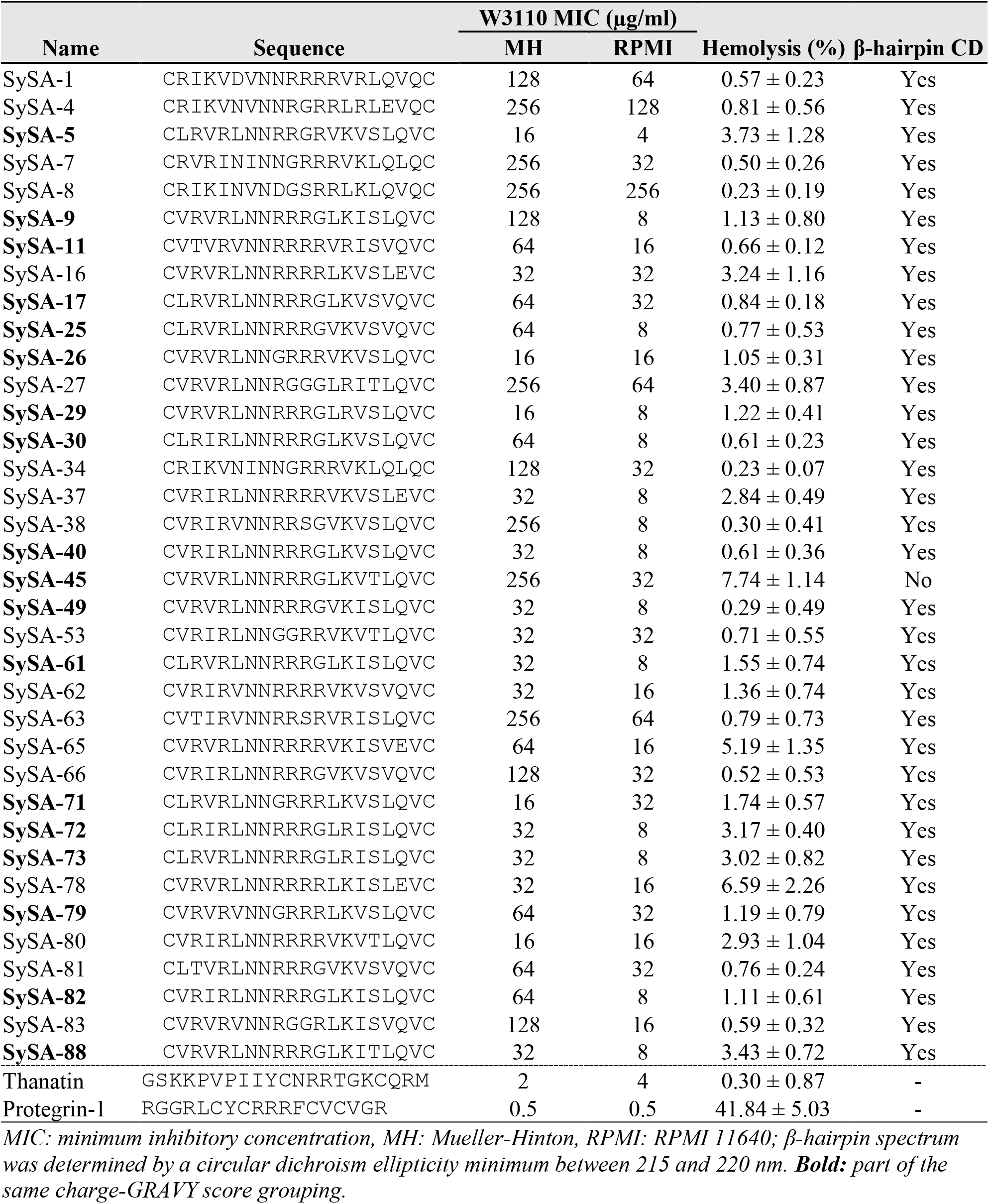
Properties of SynCH peptides verified as antibacterial.

We were curious whether SySA peptides functioned via membrane disruption like Protegrin-1 or by a periplasmic mode of action like Thanatin (*11*). To investigate, we measured the amount of propidium iodide (PI) uptake caused by our select SySA peptides and compared them to Protegrin-1 and Thanatin (**Fig. 3B**). PI fluorescence occurs upon DNA binding indicating that both the outer and inner membrane have been permeabilized. We treated *E. coli* W3110 cells with two-fold serial dilutions of peptide and observed PI uptake by relative fluorescence units (RFUs). All five or our SySA peptides showed high levels of PI uptake, similar to Protegrin-1, though requiring higher concentrations (**Fig. 3B**). As expected, Thanatin showed no PI uptake indicating it is not inner membrane disruptive.

Most natural β-AMPs like Protegrin-1 are also highly toxic to mammalian cells because they lack membrane specificity. For this reason, we examined each SySA peptide for its toxicity using a hemolysis assay measured as a percentage of red blood cells lysed by 300 μg/ml peptide compared to full lysis with a detergent (**Table 2**). Remarkably, our select SySA showed negligible hemolysis (< 3.73%) with some, like SySA-49, containing virtually no hemolysis (0.29 ± 0.49%) (**table S2, Fig. 3C**). For comparison, Protegrin-1 lysed 41.8 ± 5.03% of red blood cells and Thanatin, which does not function via membrane disruption, lysed only 0.3 ± 0.87% of red blood cells (**table S2, Fig. 3C**). This result suggests that active SySA peptides function through membrane disruption but demonstrate bacterial membrane selectively. This is an uncommon characteristic for members of the β-AMP class.

Lastly, we questioned if the naïve peptide sequences discovered through our screen could be optimized to improve their activity. Results from our biochemical characterization suggest high charge is important. Previous data from optimization of another SLAY identified β-AMP found that shorter peptide length and additional disulfide bonds increased potency (*25*). For these reasons we generated a 27-peptide optimization library around our most potent peptide (SySA-5) by shortening its length, increasing its charge, and adding the potential for a second intramolecular disulfide bond while also maintaining alternating residue side chain properties in the antiparallel β-sheets (**table S1**).

We had this library commercially synthesized and tested its ability to inhibit the growth of *Acinetobacter baumannii* AB5075, a clinically isolated Carbapenem Resistant Acinetobacter (CRA) pathogen (*28*). MICs were performed in MH as well as 100% fetal bovine serum (FBS), which is likely more representative of the *in vivo* environment. We found that 23 or our 27 variants increased potency as much as 8-fold in MH. Additionally, eleven gained the ability to kill *A. baumanii* AB5075 in FBS (**table S1**). This was surprising as degradation in serum is a common limitation of peptide use. Curious whether increased potency correlated with increased toxicity we also measured each variants hemolysis at 128 μg/ml and reported it as a fold change relative to SySA-5 (**table S1**). Most had less than a two-fold increase relative to SySA-5. The most potent of the variants (SySA 5.17) had the same MIC against *A. baumanii* AB5075 in FBS as Protegrin-1 (32 μg/ml), but was greater than ten-fold less hemolytic at 128 μg/ml. This data suggests the SynCH peptides can be easily optimized to have greater therapeutic potential than naturally occurring β-AMPs.

### SySA peptides have a constrained cyclic β-hairpin structure

Natural antibacterial peptides, including β-AMPs, generally require membrane or membrane mimics like lipopolysaccharide (LPS) to form alpha helical or β-hairpin secondary structures (*24*, *25*). For example, our natural β-hairpin peptide controls have a single molar ellipticity minimum at 200 nm in phosphate buffer alone, consistent with a random coil secondary structure **(Fig. 3D**) (*23*); however, the CD spectra of all but one of the 36 active SySA peptides (SySA-45) have a single minimum between 215 and 220 nm consistent with a β-hairpin secondary structure (**Table 2, Fig. 3D, and fig. S3**). This suggests that SySA peptides are more conformationally constrained than many natural β-AMPs. However, this alone does not dictate antibacterial activity because most of our inactive SySA peptides are also conformationally constrained (**fig. S3**). Conformational changes in β-AMP structure upon membrane interaction have been implicated in causing mammalian cytotoxicity (*29*). This could help explain the low hemolysis observed with our SynCH peptides.

Next, we looked to confirm that our most active SySA peptides were also cyclized via a disulfide bond by using LC/MS analysis. All five of the most potent SySA peptides also had a majority of their molecular population participating in a disulfide bond (**Fig. 3E**). Three of these peptides (SySA-5, 26, and 29) were 100% cyclized in solution. This data, combined with our CD analysis, strongly suggests that active SySA peptides have a macrocyclic β-hairpin structure which is conformationally constricted, a unique feature for members of the β-AMP class.

### Machine learning identifies features important for antibacterial activity

The design of the SynCH library reliably produced cyclized β-hairpins with low/no hemolytic activity, but identifying sequences features responsible for antibacterial activity was more challenging. To overcome this hurdle, we trained a machine learning algorithm to predict peptide potency with 80% of our SySA biochemical data and used the remaining 20% for validation. For a detailed description of our model please see the supplemental text. The final trained model was able to explain 90% of the variation (*R*^2^ = 0.90, *p* < 2.2 × 10^-16^) in the training dataset and 78% of the variation (*R*^2^ = 0.78, *p* = 2.5 × 10^-8^) in the validation dataset. This implies the trained model accurately predicted the activities of SySA sequences set aside for validation (**Fig. 4A**). A predictive potency score was then generated using this algorithm for all 196,608 peptide sequences in the SynCH library.

**Figure 4.**
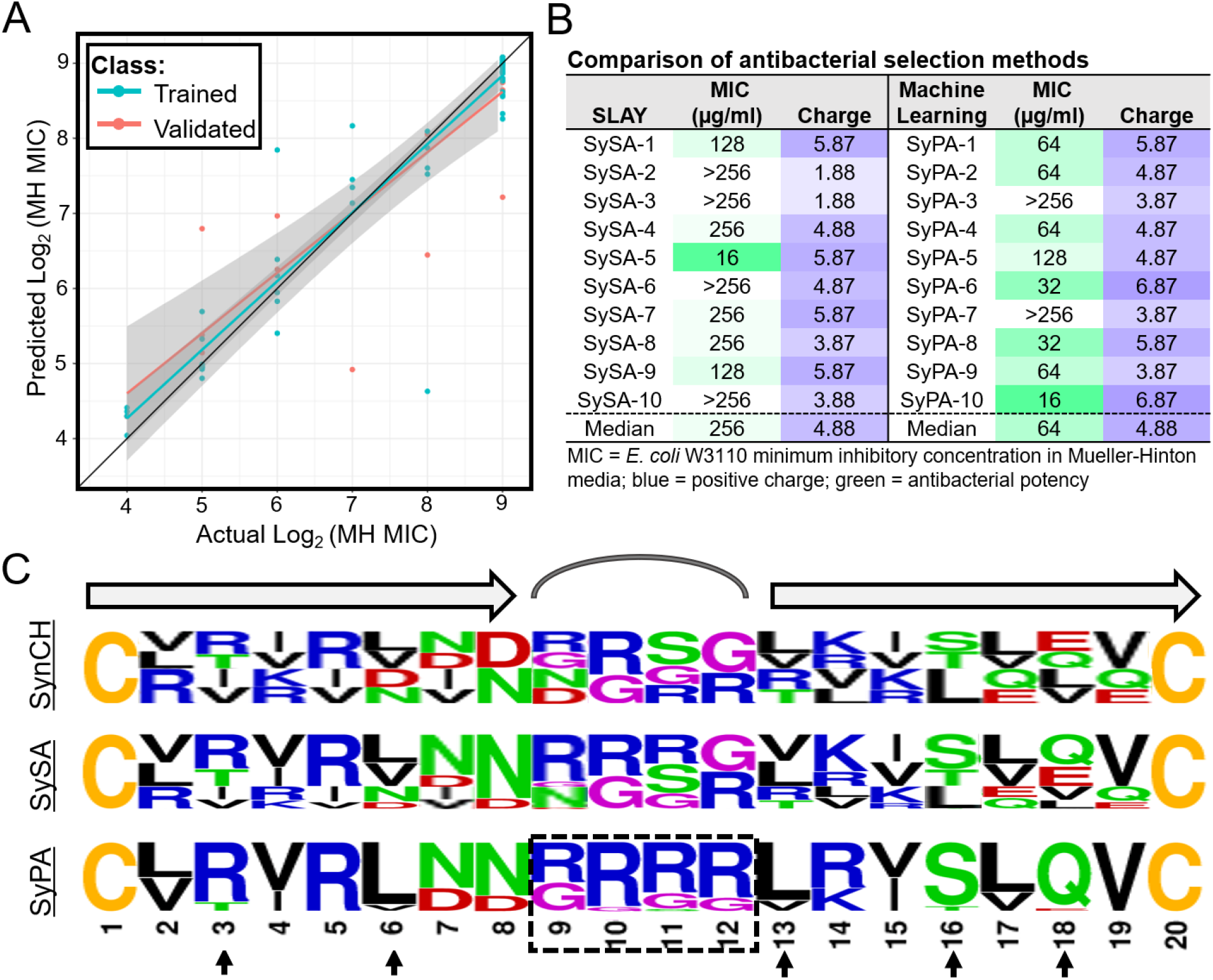
Machine leaning identifies features important for activity. (**A**) Actual versus predicted log_2_ MIC models produced via training and validation of the machine learning algorithm. (**B**) Table listing the MIC and charge of peptides identified with SLAY (left panel), or the machine learning (ML) (right panel). (**C**) residue frequencies at each position of 100 SynCH peptides (top), the top 100 SySA peptides (middle) and the top 100 SyPA peptides (bottom). Residues are color coded based on side chain properties: blue = basic, red = acidic, green = polar, black = aliphatic, purple = no side chain, yellow = sulfur containing.

To assess the accuracy of our predictive modeling we randomly selected 20,000 SynCH peptides excluded from our SLAY analysis and chose the ten with the lowest predicted potency score (SynCH Predicted Active, or “SyPA”). We had these ten peptides commercially synthesized and measured their MIC in MH against *E. coli* W3110. Remarkably, 8/10 SyPA peptides were active with MIC values ranging from 16 to 128 μg/ml (median MIC 64 μg/ml, **Fig. 4B**). This was a vast improvement to our top ten SLAY screening results where only 6/10 peptides were active and less potent overall (median MIC 256 μg/ml, **Fig. 4B**). One predicted peptide (SyPA-10) was as potent in MH media as any of the top 81 SySA peptides we examined. Furthermore, our modelling also identified active peptides with charges less than 5.8 (SyPA-2, 4, 5, and 9) and these peptides were more potent than peptides of similar charge identified using SLAY.

To explore the sequence features most important for predicting antibacterial potency, we plotted the residue frequency at each position for 100 random SynCH peptides and compared them to the top 100 peptides selected by SLAY (SySA) and predicted by machine learning (SyPA) (**Fig. 4C**) The SySA peptides had few obvious differences from SynCH other than a slight preference for the aliphatic first library. However, several sequence features were clearly enriched in the SyPA group (**Fig. 4C,**bottom). SyPA peptides were almost exclusively from the aliphatic first library possibly because of a greater potential for arginine in their loop region, which was also clearly enriched. Arginine was also highly enriched at position three. Additionally, leucine was highly preferred over valine at positions six and thirteen. Lastly, serine and glutamine were vastly favored over threonine and glutamate at positions sixteen and eighteen, respectively.

## Discussion

The goal of peptide design is to develop *de novo* sequences with and intended structure and activity. Here, we successfully designed and produced a large peptide library with predominantly macrocyclic β-hairpin structure and identified those with antibiotic activity. Our design lacked several features commonly used in β-hairpin peptide design including the use of proline in the loop region and aromatic residues like tryptophan in the β-sheet regions (*1*). In contrast, arginine was highly tolerated in both the loop region and β-sheets. We found this remarkable, as the loop sequence is considered critical for final β-hairpin stability (*30*), yet arginine is not often considered in loop design.

The β-AMPs identified from our library killed through membrane disruption but were not hemolytic. This membrane selectivity is highly unusual for β-AMPs (*13*) and warrants further study. It is possible that the constrained structure of SynCH peptides helps contribute to this selectivity. Conformational change has been shown to increase cytotoxicity in variants of Protegrin-1 (*29*). However, the lack of aromatic residues in SynCH peptides could also contribute.

Our machine learning algorithm accurately predicted active SynCH peptides with greater accuracy than our cell-based approach and helped identify several sequence features associated with antibacterial potency. Most prominent was the enrichment of arginine, especially within the loop region which could also explain a preference for the aliphatic first library. This enrichment increases the overall charge of the peptide and is likely important for interaction with the negatively charged outer membrane. The reasons for the other enriched residues are less clear, but these preferences could be related to preferred inter-strand contacts between the antiparallel β-sheets.

We are excited about the future possibilities of pairing functional cell-based peptide screening technology with machine learning strategies, especially for antibiotic discovery. We believe as more antibacterial peptide data becomes available through synthetic screening technology, machine learning may be able to predict antibacterial activity *de novo,* bypassing the need for human design and cell-based screens entirely.

## Acknowledgments

We would like to thank the Targeted Therapeutic Drug Discovery and Development Program for access to circular dichroism training and equipment. Also, Dr. Kristin Blake at the Mass Spectrometry Facility at The University of Texas at Austin for her help with high-resolution LC/MS.

## Author contributions

Conceptualization: JR, CD, BW, TC, CW Methodology: JR, CD, TC,

Investigation: JR, CD, GD, TC, KG

Visualization: JR, CW

Funding acquisition: BW, CW

Project administration: JR, BW

Supervision: JR, BD, CW

Writing – original draft: JR

Writing – review & editing: JR, BW, CW

## Data and materials availability

Any data not available in the manuscript or supplementary material can be made available upon request to the corresponding author.

## Supplementary Materials

Contains the Materials and Methods, Supplemental Text, Figures S1-S3, and Table S1-S2

