## Supplemental Materials for "Designing β-hairpin peptide macrocycles for antibiotic potential"

### Supplementary Materials

Justin R. Randall, Cory D. DuPai, T. Jeffrey Cole, Gillian Davidson, Kyra E. Groover, Claus O. Wilke, and Bryan W. Davies

**This PDF file includes:**

Materials and Methods
Supplementary Text
Figs. S1 to S3
Tables S1 to S2

### Materials and Methods

SynCH library cloning

Detailed methods for library creation have been previously reported (*26*, *27*). Briefly, the two library inserts were generated by PCR using forward primer oJR557 with reverse primers oJR560 and oJR561 encoding each library. The 2x(NR)tether gBlock was used as the template. Both inserts and the pMMBEH67_lpp_ompA vector were digested with KpnI and SalI independently and the two libraries were ligated overnight at 4°C using T4 ligase. The two ligated libraries were cleaned and transformed into *E. coli* 10-β competent cells (NEB) and plated on large bioassay dishes (250 mm) filled with LB agar supplemented with 75 μg/ml carbenicillin and grown overnight. The next morning colonies were counted, scraped, pooled, aliquoted, and frozen in 10% glycerol for long term storage. One ml of pooled cells was thawed, and maxi prepped to obtain the combined SynCH plasmid library. This library was then transformed into *E. coli* W3110 competent cells for further analysis via SLAY (see below).

Peptide synthesis

All peptides used in this work were synthesized commercially by GenScript's custom peptide synthesis service. Each peptide goes through reverse phase high performance liquid chromatography (RP-HPLC) and mass spectrometry quality control analysis to confirm purity and molecular weight. Lyophilized peptides were resuspended in water at 10 μg/ml. A full list of all of the peptides used and their purity can be found in supplemental excel file 1.

Circular dichroism spectroscopy

Stock peptides were diluted in phosphate buffered saline (pH 7.4) to 200 μg/ml in a volume of 200 μl. Samples were incubated 1-2 hours at room temperature and then analyzed using a Jasco-815 CD spectrometer with a 0.1 cm path-length quartz cuvette at the Targeted Therapeutic Drug Discovery & Development Program Core at UT Austin. The CD spectra were collected using far-UV spectra (195-250 nm) with background corrected for phosphate buffered saline alone. Ellipticity was converted from mdeg to molar ellipticity. Reported spectra are an average of three separate spectra obtained from the same sample and adjusted for molar concentration.

High resolution mass spectrometry

Stock peptides were diluted into phosphate buffered saline (pH 7.4) at 0.1 mg/ml in a volume of 150 μl and placed into 2 ml autosampler vials with small volume inserts (Fisherbrand 03-391-8). Samples were separated by a C8 liquid chromatography column, and an Extracted Ion Chromatogram (EIC) was generated for the most prevalent charge state isotopes. Mass spectra were generated for each extracted LC peak using an Agilent Technologies 6546 Accurate-Mass Q-TOF LC/MS instrument. Analysis was performed using Agilent MassHunter Qualitative software v10. The isotope distribution for each LC peak from each sample was compared to predicted distributions created using Agilent's Isotope Distribution Calculator. Percentages of molecules with disulfide bonds was determined by comparing the area under the curve for each LC peak containing a disulfide bond over the total area of all relevant peaks (Sum area %) and reported as a mean of three technical replicates ± one standard deviation.

Minimum inhibitory concentration

MICs were performed by a modified CSLI procedure. Stock peptides were diluted to 2.56 μg/ml in 50 μls of 0.01% acetic acid containing 0.2% bovine serum albumin (BSA). Peptide was then serially diluted two-fold in this solution and 10 µl of each dilution was added to a polypropylene 96 well plate (Corning REF# 3879) in triplicate. Separately, *E. coli* W3110 or *A. baumannii* AB5075 was grown overnight in 5 ml MH at 37 °C. Cells from these cultures were diluted to a concentration of 5-6 x $10^5$ cells/ml in either MH, RPMI, or FBS and 90 µl were added to each well of the 96 well plate containing diluted peptide. Plates were wrapped twice in parafilm and incubated at 37 °C for 18-24 hours. Wells were examined by eye against wells with no bacteria for signs of growth. MICs were reported as the minimum concentration with no observable growth. In cases where triplicate samples differed, the concentration supported by the median of the three replicates was reported.

<u>Surface localized antimicrobial display procedure</u>

SLAY procedures have been detailed previously (*26, 27*). Briefly, 100 µl of *E. coli* W3110 frozen cells containing the pooled SynCH plasmid library were recovered in 10 ml of LB supplemented with 75 µg/ml carbenicillin for 2 hours. The culture was then back diluted to OD 0.05 and three triplicate 5 ml cultures were set up in LB supplemented with 75 µg/ml carbenicillin. Triplicate reactions included: Uninduced (0 uM IPTG), Low induction (15 uM IPTG), and High induction (100 uM IPTG). All triplicate cultures were then grown for 3 hours at 37°C. Plasmids from each triplicate culture were isolated via miniprep and Illumina sequencing primers (See table S4) were used to produce an amplicon via PCR from plasmids from each culture containing a unique i7 barcode identifier. Amplicons were then gel extracted, cleaned, and concentrated, and sent to Genewiz for next generation sequencing by Illumina HiSeq technology with 30% added Phi-X DNA.

<u>SLAY sequencing analysis.</u>

Methods of SLAY analysis have been previously described elsewhere (*26*, *27*). Here, raw sequencing reads were trimmed of adapter sequences using flexbar (31) and then assessed for quality via FastQC (*32*). Next, trimmed reads were mapped to a reference derived from all possible SynCH sequences using Kallisto (*33*). Finally, $\log_2$fold change estimates and other relevant statistics were calculated using DESeq2 (*34*).

<u>Hemolysis</u>

Single donor human red blood cells (Innovative Research #IWB3ALS) were washed in PBS and adjusted to a concentration of 1 x $10^9$ cells/ml. Each peptide was added individually to 200 µl of cells at a concentration of either 128 or 300 µg/ml in a 96 well polypropylene plate (Corning 3879). PBS alone and 1% triton-X100 were used for background normalization and 100% hemolysis respectively. Each reaction was set up in triplicate. Plates were incubated for 3 hours at 37 ºC. Following incubation samples were centrifuged at 800 g for 20 min and 100 µl of supernatant was transferred to a flat bottom 96 well plate (Genesse 25-104). Percent hemolysis for each sample was determined by normalizing the absorbance at 540 nm for each sample to the average background and dividing by the average absorbance for 1% Triton X-100 (100% hemolysis). Error bars represent one standard deviation of triplicate samples.

<u>Propidium iodide uptake</u>

Propidium iodide uptake was measured for *E. coli* W3110 using the as previously described (*25*). Briefly, single colonies were from overnight growth on LB were inoculated into Mueller Hinton broth and grown to mid-log phase.  Cells were then washed twice with 1X PBS + 50mM glucose and resuspended to a final optical density of 0.1 in 1x PBS + 50mM glucose. Propidium iodide was added to the cells at a concentration of 10 µg/mL (15 µM) and 50 µL of the propidium iodide cell mixture was quickly added to a pre-prepared 96-well plate (NUNC black-walled clear bottom plate) that contained 50 µL of different peptide concentrations. The plate had been prepared by serially diluting peptides two-fold in 1x PBS + 50 mM glucose, the highest concentration of peptides was equivalent to 128 µg/mL, which with the addition of cells became 64 µg/mL. The plate was then allowed to incubate for 25 minutes in the dark at 37°C. After, 25 minutes had elapsed the plate was read using the Biotek Synergy LX plate reader with the fluorescence red filter cube* every 5 minutes for 25 minutes with shaking in between. To account for background fluorescence each plate contained three triplicate columns with non-treated cells in 1X PBS + 50mM glucose. Triplicate samples were normalized to non-treated wells and presented as a mean ± one standard deviation (n=3).

Machine learning and regression modelling

Machine learning models were trained to predict MIC and hemolysis from peptide sequences. For the MIC model, values were first $\log_2$ transformed. Peptide sequences were embedded as numerical vectors using the Bepler deep protein language model (*35*) from the bio-embeddings python library. The peptide sequences were split into a training set that represented a random sample 80% the size of the original dataset and the remaining 20% was set aside as the test dataset. An automated machine learning (autoML) approach (*36*) was used to find the best machine learning regression model that minimized the Root Mean Squared Error (RMSE). The mljar library was used to carry out the autoML model selection with varying combinations of machine learning architecture, hyperparameters, and feature selection. The combination of model architecture, hyperparameters, and feature selection that minimized RMSE was chosen as the final model.

### Supplementary Text

Machine learning algorithm and modelling

For this model, MIC values obtained for the top 81 SySA peptides were $\log_2$ transformed and amino acid sequences were embedded as numerical vectors using the Bepler deep protein language model (*34*) from the bio-embeddings python library (*38*). We then used AutoML (*36*) to fit an array of different predictors to 80% of the SySA biochemical data (the training data) and validated performance on the remaining 20% (validation data). Each fitted predictor takes a numerical embedding of a peptide as input and produces a score as output, where a lower score indicates a higher likelihood of antibacterial activity. The AutoML algorithm considered predictors such as linear, random forest, neural networks, or gradient boosting.
The best performing predictor found by AutoML was of type LightGMB (*37*), with 16 selected features. Because the features are embedding scores, we could not interpret them directly. To nonetheless gain some insight into these features, we correlated them with basic properties of peptides such as the proportion of different amino acids, the peptide length, charge, hydrophobicity, and so on. We found that three features perfectly represented the fractions of alanine, proline, and glutamine in a peptide. These and other features did not have a straightforward interpretation or obvious correlations with our library design. Correlations with

peptide length, weight, or the amount of cysteine were also common; however, these correlations had relatively low $R^2$ values of between 10% and 38%. These observations highlight that the protein embedding scores capture non-trivial aspects of peptide biochemistry that don't necessarily correspond to simple and straightforward biochemical quantities (**Fig. 4A**).

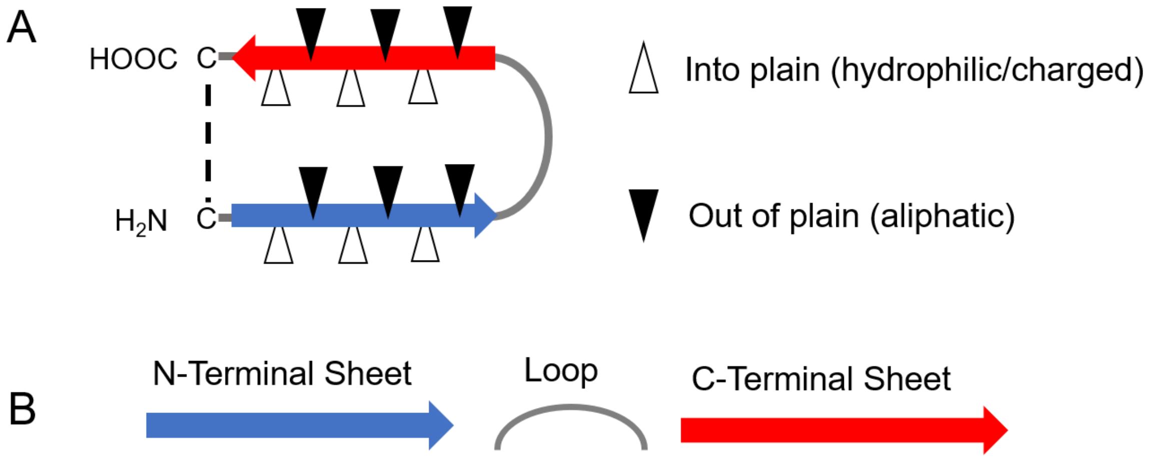


**ALIPHATIC FIRST**

| | | | | | | | | | | | | | | | | | | | | | | |
|---|---|---|---|---|---|---|---|---|---|---|---|---|---|---|---|---|---|---|---|---|---|---|
| | C | VL | RT | VI | R | VL | DN | DN | GR | GR | GSR | GR | VL | RK | VI | ST | VL | QE | V | C | 98304 | Total |
| | | | + | | + | | - | - | + | + | + | + | | + | | | | - | | | | |
| | | | 0.5 | | 1 | | -0.5 | -0.5 | 0.5 | 0.5 | 0.34 | 0.5 | | 1 | | | | -0.5 | | | 2.84 | Charge |
| **CODONS** | **TGT** | **STT** | **ASG** | **RTT** | **AGA** | **STT** | **RAT** | **RAT** | **SGA** | **SGA** | **VGT** | **SGT** | **STT** | **ARA** | **RTT** | **AST** | **STT** | **SAA** | **GTG** | **TGT** | | |
| | | CTT | ACG | ATT | | CTT | AAT | AAT | CGA | CGA | CGA | CGT | CTT | AAA | ATT | ACT | CTT | CAA | | | | |
| | | GTT | AGG | GTT | | GTT | GAT | GAT | GGA | GGA | GGA | GGT | GTT | AGA | GTT | AGT | GTT | GAA | | | | |
| | | | | | | | | | | | AGA | | | | | | | | | | | |

**CHARGED FIRST**

| | | | | | | | | | | | | | | | | | | | | | | |
|---|---|---|---|---|---|---|---|---|---|---|---|---|---|---|---|---|---|---|---|---|---|---|
| | C | R | IV | RK | VI | DN | VI | DN | DN | GR | GSR | GR | TR | LV | RK | L | QE | LV | QE | C | 98304 | Total |
| | | + | | + | | - | | - | - | + | + | + | + | | + | | - | | - | | | |
| | | 1 | | 1 | | -0.5 | | -0.5 | -0.5 | 0.5 | 0.34 | 0.5 | 0.5 | | 1 | | -0.5 | | -0.5 | | 2.34 | Charge |
| **CODONS** | **TGT** | **AGA** | **RTT** | **ARA** | **RTT** | **RAT** | **RTT** | **RAT** | **RAT** | **SGA** | **VGC** | **SGT** | **ASG** | **STT** | **ARA** | **CTG** | **SAA** | **STT** | **SAA** | **TGT** | | |
| | | | ATT | AAA | ATT | AAT | ATT | AAT | AAT | CGA | CGA | CGT | ACG | CTT | AAA | | CAA | CTT | CAA | | | |
| | | | GTT | AGA | GTT | GAT | GTT | GAT | GAT | GGA | GGA | GGT | AGG | GTT | AGA | | GAA | GTT | GAA | | | |
| | | | | | | | | | | | AGA | | | | | | | | | | | |

**196608 Combined**

**Fig. S1. High-resolution mass spectrometry of cyclized symbah-1, Related to Figure 1.** (**A**) Diagram of our basic β-hairpin peptide design. (**B**) Aliphatic first and charged first SynCH libraries showing the separate regions and each potential amino acid at every position. The average charge contribution at each position is shown and the codon variation used to potentiate the different amino acids at each position are below.

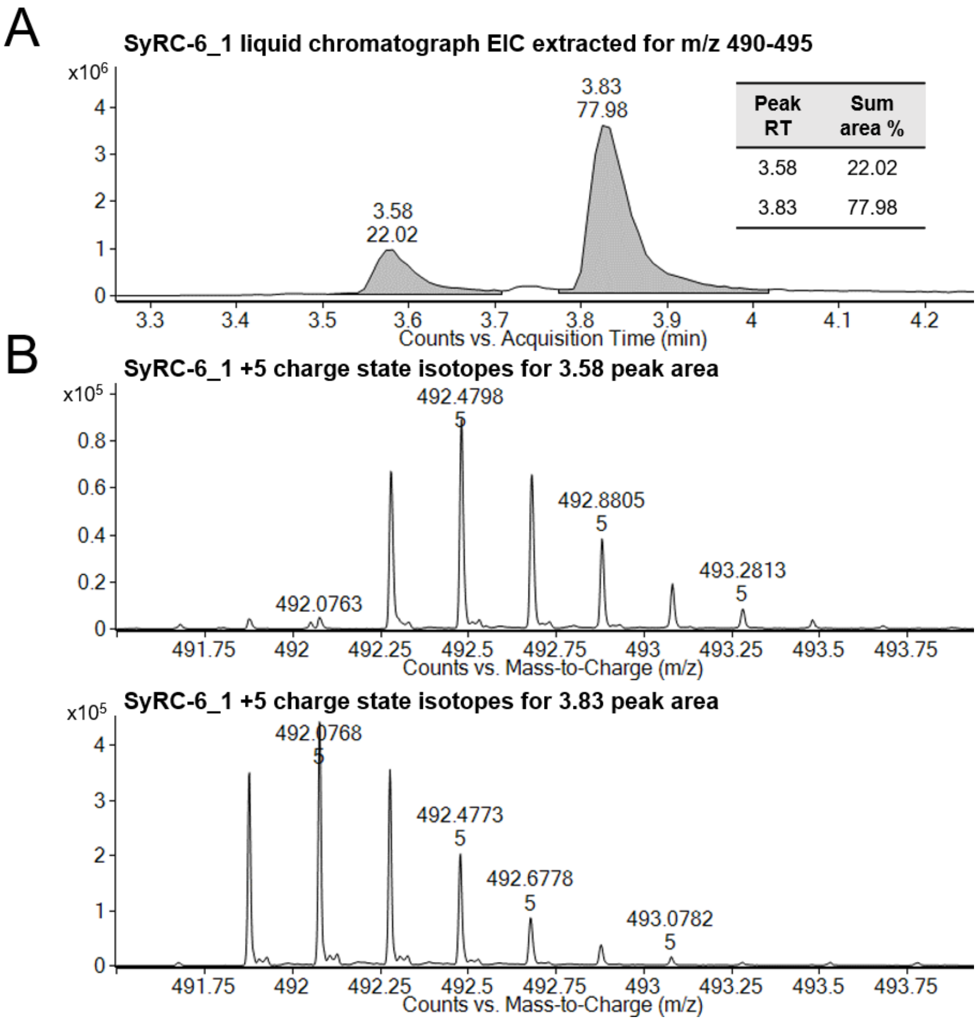


**Fig. S2. Calculating peptide percent cyclization, Related to Figure 3E and table S1.**
(**A**) Liquid chromatography elution peaks from SyRC-6 in PBS. Peak acquisition time and percentage of total elution peaks are in the inset table. (**B**) +5 charge state peaks with the measured monoisotopic mass ($M_{mi}$) of each peak. Data here represents the measurement of a single sample. the ~2 Dalton drop in monoisotopic mass between the two peaks corresponds to the formation of a disulfide bond.

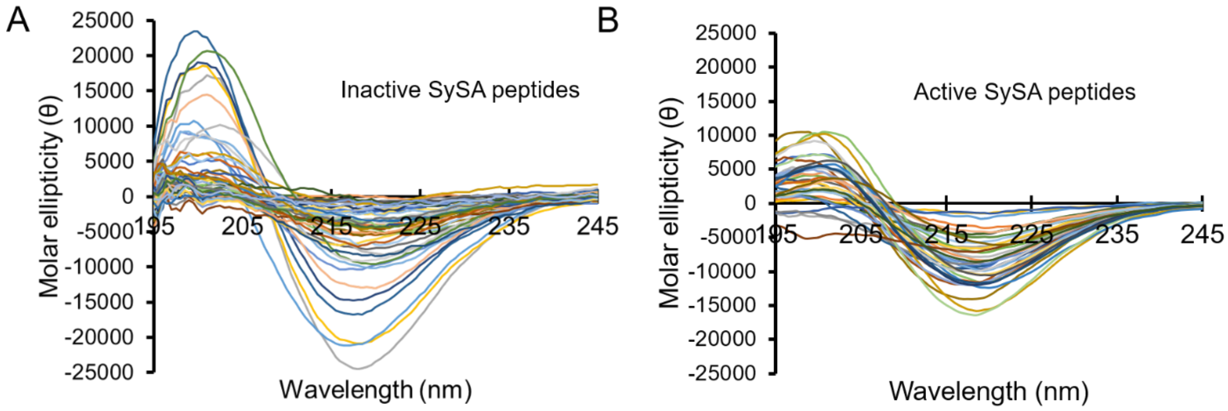


**Fig. S3. SynCH SLAY active peptide structures, Related to Figure 3D and table S2.** Circular dichroism spectra of the inactive (**A**) and active (**B**) top 88 SySA peptides. Spectra with a single Molar ellipticity minimum between 215-220 was considered to contain a β-hairpin secondary structure.

**Table S1. SySA-5 *A. baumannii* AB5075 optimization library, Related to Figure 3.**

| Name | Sequence | AB5075 MIC (µg/ml) MH | AB5075 MIC (µg/ml) FBS | Length (AAs) | Potential Cysteine pairs | # Cationic AAs | Fold Change Hemolysis |
|---|---|---|---|---|---|---|---|
| SySA-5 | CLRVRLNNRRGRVKVSLQVC | 32 | >128 | 20 | 1 | 6 | 1.0 |
| SySA-5.1 | CLRVRL**R**NRRGRVKVSL**R**VC | 8 | 128 | 20 | 1 | 8 | 1.8 |
| SySA-5.2 | CLRVRL**R**NRR**R**RVKV**R**L**R**VC | 4 | >128 | 20 | 1 | 10 | 2.6 |
| SySA-5.3 | CLRVRL**C**NRR**RC**VKVRL**R**VC | 8 | 128 | 20 | 2 | 8 | 1.8 |
| SySA-5.4 | CLRVRL**CR**RR**RC**VKV**R**L**R**VC | 8 | 64 | 20 | 2 | 10 | 2.9 |
| SySA-5.5 | C-RVRLNNRRGRVKVSLQ-C | 8 | >128 | 18 | 1 | 6 | 0.9 |
| SySA-5.6 | C-RVRL**R**NRRGRVKVSL**R-**C | 4 | 128 | 18 | 1 | 8 | 0.7 |
| SySA-5.7 | C-RVRL**R**NRR**R**RVKV**R**L**R-**C | 8 | >128 | 18 | 1 | 10 | 0.9 |
| SySA-5.8 | C-RVRL**C**NRR**RC**VKVSL**R-**C | 8 | >128 | 18 | 2 | 7 | 0.8 |
| SySA-5.9 | C-RVRL**CR**RR**RC**VKV**R**L**R-**C | 8 | >128 | 18 | 2 | 9 | 0.6 |
| SySA-5.10 | CLRVRLN-RRGR-KVSLQVC | 4 | 128 | 18 | 1 | 6 | 1.4 |
| SySA-5.11 | CLRVRL**R-**RRGR**-**KV**R**LQVC | 4 | 64 | 18 | 1 | 8 | 3.0 |
| SySA-5.12 | CLRVRL**R-**RR**R**R-KV**R**L**R**VC | 4 | >128 | 18 | 1 | 10 | 3.7 |
| SySA-5.13 | CLRVRL**C-**RRGR-**C**VSL**R**VC | 16 | >128 | 18 | 2 | 6 | 3.2 |
| SySA-5.14 | CLRVRL**C-**RR**R**R-**C**V**R**L**R**VC | 4 | 64 | 18 | 2 | 8 | 5.6 |
| SySA-5.15 | C-RVRLN**C**RRGR**C**KVSLQ-C | 16 | >128 | 18 | 2 | 6 | 1.3 |
| SySA-5.16 | C-RVRL**RC**RRGR**C**KVSL**R-**C | 32 | >128 | 18 | 2 | 8 | 1.3 |
| SySA-5.17 | C**-**RVRL**RC**RR**R**R**C**KV**R**L**R-**C | 4 | 32 | 18 | 2 | 10 | 2.4 |
| SySA-5.18 | CLRVRLN-RR-R-KVSLQVC | 8 | 128 | 17 | 1 | 6 | 1.0 |
| SySA-5.19 | CLRVRL**R-**RR-R-KV**R**LQVC | 4 | >128 | 17 | 1 | 8 | 2.5 |
| SySA-5.20 | CLRVRL**R-**RR-R-KV**R**L**R**VC | 4 | 128 | 17 | 1 | 9 | 3.3 |
| SySA-5.21 | CLRVRL**C-**RR-R-**C**VSL**R**VC | 16 | >128 | 17 | 2 | 6 | 2.4 |
| SySA-5.22 | CLRVRL**C-**RR-R-**C**V**R**L**R**VC | 32 | 64 | 17 | 2 | 7 | 2.1 |
| SySA-5.23 | C-RVRLN-RRGR-KVSLQ-C | 8 | >128 | 16 | 1 | 6 | 1.0 |
| SySA-5.24 | C-RVRL**R-**RRGR-KVSL**R-**C | 16 | >128 | 16 | 1 | 8 | 1.1 |
| SySA-5.25 | C-RVRL**R-**RR**R**R-KV**R**L**R-**C | 32 | >128 | 16 | 1 | 10 | 1.5 |
| SySA-5.26 | C-RVRL**C-**RR**R**R-**C**V**R**L**R-**C | 8 | >128 | 16 | 2 | 8 | 0.8 |
| SySA-5.27 | C-RVR**CR-**RR**R**R-K**CR**L**R-**C | 128 | >128 | 16 | 2 | 10 | 0.6 |
| Protegrin-1 | RGGRLCYCRRRFCVCVGR | 1 | 32 | 18 | 2 | 6 | 32.7 |

MIC: minimum inhibitory concentration, MH: Mueller-Hinton, FBS: fetal bovine serum, AA: amino acid; **bold:** residue change; green: fold-improvement, red: fold-deterioration

**Table S1. Plasmids, Strains, and Oligonucleotides.**

| Plasmids | Source |
|---|---|
| pMMBEH67_lpp_ompA | (*26*) |
| **Strains** | **Source** |
| *E. coli* W3110 | Lab Stock |
| *A. baumannii* AB5075 | (*28*) |
| **Oligonucleotides** | **Sequence** |
| oJR557 - F SynCH | gtattggtaccagtcaagagcctg |
| oJR560 - R SynCH Aliphatic First | ctg cag gtc gac tta ACA CAC TTS AAS AST AAY TYT AAS ACS ACB TCS<br>TCS ATY ATY AAS TCT AAY CST AAS ACA ggt tcc tcc gat acc cgc ag |
| oJR561 - R SynCH Charged First | ctg cag gtc gac tta ACA TTS AAS TTS CAG TYT AAS CST ACS GCB TCS<br>ATY ATY AAY ATY AAY TYT AAY TCT ACA ggt tcc tcc gat acc cgc ag |
| 2x(NR)tether gBlock | ATTGCCGATGGTACACGTCAAGTCAAGAGCCTGCAGCGCCCGCCGC<br>AGAGGCGACTCCTGCTGCTGAAGCTCCAGCTAGCGAAGCGCCTGCA<br>GCAGAAGCTGCCCCAGCGGATGCTGCCGAAGCCCCAGCCGCTGGCA<br>TCAGTCAGGAACCTGCTGCACCAGCTGCGGAAGCTACACCAGCAGC<br>GGAGGCACCAGCGAGTGAAGCACCGGCTGCGGAAGCCGCTCCTGCA<br>GATGCCGCTGAGGCTCCAGCTGCGGGTATCGGAGGAACCCGCGGTG<br>GGCGTCTTTGTTA |
| F amplicon | aatgATACGGCGACCACCGAGATCTACACTCTTTCCCTACACGACGCT<br>CTTCCGATCTCTCCAGCTGCGGGTATCGGAGGA |
| R index 1 | CAAGCAGAAGACGGCATACGAGATCGTGATGTGACTGGAGTTCAGA<br>CGTGTGCTCTTCCGATCTgccaagcttgcatgcctgcaggtcgacTTA |
| R index 2 | CAAGCAGAAGACGGCATACGAGATACATCGGTGACTGGAGTTCAGA<br>CGTGTGCTCTTCCGATCTgccaagcttgcatgcctgcaggtcgacTTA |
| R index 3 | CAAGCAGAAGACGGCATACGAGATGCCTAAGTGACTGGAGTTCAGA<br>CGTGTGCTCTTCCGATCTgccaagcttgcatgcctgcaggtcgacTTA |
| R index 4 | CAAGCAGAAGACGGCATACGAGATTGGTCAGTGACTGGAGTTCAGA<br>CGTGTGCTCTTCCGATCTgccaagcttgcatgcctgcaggtcgacTTA |
| R index 5 | CAAGCAGAAGACGGCATACGAGATCACTGTGTGACTGGAGTTCAGA<br>CGTGTGCTCTTCCGATCTgccaagcttgcatgcctgcaggtcgacTTA |
| R index 6 | CAAGCAGAAGACGGCATACGAGATATTGGCGTGACTGGAGTTCAGA<br>CGTGTGCTCTTCCGATCTgccaagcttgcatgcctgcaggtcgacTTA |
| R index 7 | CAAGCAGAAGACGGCATACGAGATGATCTGGTGACTGGAGTTCAGA<br>CGTGTGCTCTTCCGATCTgccaagcttgcatgcctgcaggtcgacTTA |
| R index 8 | CAAGCAGAAGACGGCATACGAGATTCAAGTGTGACTGGAGTTCAGA<br>CGTGTGCTCTTCCGATCTgccaagcttgcatgcctgcaggtcgacTTA |
| R index 9 | CAAGCAGAAGACGGCATACGAGATCTGATCGTGACTGGAGTTCAGA<br>CGTGTGCTCTTCCGATCTgccaagcttgcatgcctgcaggtcgacTTA |
| R index 10 | CAAGCAGAAGACGGCATACGAGATAAGCTAGTGACTGGAGTTCAGA<br>CGTGTGCTCTTCCGATCTgccaagcttgcatgcctgcaggtcgacTTA |
| R index 11 | CAAGCAGAAGACGGCATACGAGATGTAGCCGTGACTGGAGTTCAGA<br>CGTGTGCTCTTCCGATCTgccaagcttgcatgcctgcaggtcgacTTA |
| R index 12 | CAAGCAGAAGACGGCATACGAGATTACAAGGTGACTGGAGTTCAGA<br>CGTGTGCTCTTCCGATCTgccaagcttgcatgcctgcaggtcgacTTA |

All oligonucleotides were ordered from Integrated DNA technologies (IDT). IDT single letter nucleotide base notations are used.
